# Functional auxin signaling and phosphorus acquisition triggered in planta by the extremophile bacterium *Pseudomonas extremaustralis*

**DOI:** 10.1101/2025.08.03.668350

**Authors:** Carola Agranatti, Jose G Ibarra, Mariana B Galeano, R Terranova, Manuel S. Godoy, Nancy I López, Martiniano M Ricardi, Paula M Tribelli

**Affiliations:** Departamento de Química Biológica, Facultad de Ciencias Exactas y Naturales, Universidad de Buenos Aires, Buenos Aires, Argentina; IQUIBICEN-CONICET, Buenos Aires, Argentina; Warwick Integrative Synthetic Biology Centre (WISB), School of Life Sciences, University of Warwick, CV4 7AL UK; Departamento de Fisiología, Biología Molecular y Celular, Facultad de Ciencias Exactas y Naturales, Universidad de Buenos Aires, Buenos Aires, Argentina; IFIBYNE (UBA-CONICET), FBMC, FCEyN-UBA, Buenos Aires, Argentina

## Abstract

*Pseudomonas extremaustralis* 14-3b, a strain isolated from Antarctica, was evaluated for plant-growth-promoting traits under different oxygen conditions. Genomic analysis revealed genes related to phosphorus solubilization and indole-3-acetic acid (IAA) biosynthesis. In vitro assays confirmed its ability to solubilize tricalcium phosphate and produce gluconic acid. IAA production was higher under microaerobic conditions compared to aerobic cultures. Using *Arabidopsis thaliana* DII-VENUS reporter lines, we confirmed in vivo IAA perception in roots, with reduced VENUS fluorescence upon bacterial inoculation, regardless of prior oxygen conditions. Confocal microscopy showed root surface colonization and internal localization in cotyledons, including sub-stomatal spaces. Morphological changes in *A. thaliana* included reduced primary root length and increased root hair length. In wheat, *P. extremaustralis* exhibited chemotaxis toward root exudates and colonized root tissues. After 30 days in pot experiments, inoculated wheat plants showed significantly higher stem dry weight and leaf phosphorus content than control plants. These findings indicate that *P. extremaustralis* promotes plant growth through multiple mechanisms, supporting its potential as a bioinoculant under variable environmental conditions, including low oxygen availability.

## Introduction

The successful expansion of farmlands to increase crop production depends on the ability of plants to grow effectively in areas under suboptimal environmental conditions. The use of plant-growth-promoting rhizobacteria (PGPR) constitutes an environmentally sustainable alternative by reducing or eliminating the need for agrochemicals and enhancing crop growth, even under stressful conditions such as drought, heat, cold, limited oxygen provision (hypoxia/microaerobiosis), and high salinity (Backer et al., 2018). Plants frequently experience low oxygen conditions, particularly in tissues with high metabolic activity such as seeds, meristems, and storage organs. Oxygen deprivation leads to a metabolic shift from aerobic to anaerobic respiration, significantly reducing ATP yield. Under hypoxia, metabolic fluxes are reprogrammed, thus prioritizing essential cellular functions. Moreover, in anoxic conditions, energy production relies solely on glycolysis, causing a rapid energy deficit that can lead to cellular dysfunction and death (Pucciarello et al., 2012).

Numerous bacterial species exhibit plant growth-promoting activities, contributing to fertilization, phytostimulation, and disease control. Among the traits related to fertilization, the mobilization of phosphorus in the soil into forms suitable for plant uptake is critical, as phosphorus is one of the primary nutrients limiting plant growth (Vitousek et al., 2010). Phosphorus in soil exists in organic or inorganic forms that are not directly assimilable by plants. Organic phosphates can be mineralized by various bacterial species through the action of phosphatase, phytase, and phosphonatase enzymes (Kumar et al., 2016; Shrivastava et al., 2018 ). Inorganic insoluble phosphorus can be solubilized indirectly through the production of low-molecular-weight organic acids, mainly gluconic acid (Glick, 2012). Phytostimulation involves the production of plant growth-related phytohormones, such as indole-3-acetic acid (IAA) (Spaepen and Vanderleyden, 2011).

The expansion of agriculture into less favorable areas, combined with the effects of climate change, imposes additional environmental stress on crops. Therefore, the search for PGPR from extreme environments capable of exhibiting beneficial traits under such conditions represents a promising strategy to mitigate plant stress and enhance growth. *Pseudomonas* species are versatile bacteria with efficient colonization abilities, a wide variety of energy-generation metabolisms, and significant roles in nutrient cycles. Several species within this genus exhibit plant growth-promoting traits. *Pseudomonas extremaustralis* 14-3b is a non-pathogenic species isolated from a temporary pond in Antarctica (López et al., 2009), showing high resistance to thermal, oxidative, and nitrosative stress, and the capacity to thrive in cold environments, besides possessing genomic information to cope with osmotic stress (Ayub et al., 2004; Ayub et al., 2009; Raiger Iustman et al., 2015; Solar Venero et al., 2024). It is also able to grow under microaerobic or anaerobic conditions through nitrate reduction or arginine and pyruvate fermentation, showing, also, well-developed biofilm formation and cellular aggregation capacities (Tribelli et al., 2010; Tribelli et al., 2011; Benforte et al., 2018).

Soil represents a heterogeneous environment characterized by oxygen gradients and microenvironments, resulting from environmental conditions, bioactivities, and interactions, which are particularly intense in the rhizosphere, the zone surrounding the root (Uteau et al., 2015). Then, plant growth-promoting bacteria are exposed to oxygen gradients as they carry out their activities. Bacterial physiology is deeply impacted by oxygen tension and redox state, and the transition from aerobic to microaerobic metabolism is regulated in Pseudomonas species by the global regulator Anr, which is functional in *P. extremaustralis* (Tribelli et al., 2010; Tribelli et al., 2013; Tribelli et al., 2019).

The aim of this study was to investigate the plant-growth-promoting traits of *P. extremaustralis* under different oxygen tensions in vitro and the impact of oxygen conditions in cultures before plant inoculation using various hosts, including wheat, a commercially important crop in Argentina, and the model plant *Arabidopsis thaliana*.

## Materials and methods

### *Bacterial strains* and bacterial culture conditions

*Pseudomonas extremaustralis* 14-3b (DSM 25547) isolated in our laboratory from a sample from Antarctica (López et al., 2009; Tribelli et al., 2012) was used in this study. *P. extremaustralis* expressing GFP or mCherry (Benforte et al., 2018) were used for root colonization assays. For both aerobic and microaerobic cultures, precultures were performed in Lysogeny Broth (LB) under aerobic standard conditions and were used to inoculate bacterial cultures used for different assays. LB aerobic cultures were performed using Erlenmeyer flasks with 1:10 medium: total volume and shaking of 200 rpm, while for microaerobic cultures, sealed bottles with LB supplemented with 0.08% of KNO^3^ were used with 1:4 medium: total volume with slow shaking (50 rpm) to avoid aggregates (Tribelli et al., 2010). For all experiments, aerobic cultures were incubated for 24 h while microaerobic cultures were incubated for 48 h, both at 28°C.

### In vitro evaluation of plant-growth-promoting traits

#### Phosphate solubilization on solid medium

The ability to solubilize phosphate was determined through a plate solubilization assay using NBRIP medium plates (Nautiyal, 1999) supplemented with 5 g l^-1^ of calcium phosphate (Ca₃(PO₄)₂), 1% glucose, and 1.5 g l^-1^ of agar. Plates were inoculated with 10 µl drops of aerobic or microaerobic cultures. Phosphate solubilization was visualized as the formation of clear halos around the bacterial spots after incubation at 30°C for 7 days. The diameter of the solubilization halo and the colony diameter were measured by using Fiji (Schindelin et al., 2012).

#### Organic acid production detection

To analyze the production of organic acids, bacterial cultures were performed under aerobic conditions at 28 °C in NBRIP broth medium supplemented with tricalcium phosphate as the sole source of phosphorus. Samples of 1.5 ml were taken every 24, 48, and 72 h of culture, centrifuged, and the supernatants were frozen at -20 °C. Organic acids composition was analyzed following Scervino et al. (2011). Briefly, filtered supernatants (0.22 µm nylon filter) of NBRIP broth cultures were analyzed by high-performance liquid chromatography (HPLC), using an Aminex column HPX-87-H (Cat no. 125–0140) and a UV detector set to 215 nm at 50°C. The mobile phase consisted of 5 mmol l^-1^ H^2^SO^4^ run at a flow rate of 6 ml min^-1^. Standards of commercial organic acids were used to quantify by comparing peak areas of chromatograms with those of standards. Three independent cultures were used.

#### Indole acetic acid (IAA) determination

To evaluate IAA production, bacterial strains were cultured under aerobic and microaerobic conditions in E2 medium supplemented with 1 g·l⁻¹ tryptophan. After incubation, cultures were centrifuged at 13,000 rpm for 5 minutes, and the supernatants were collected and filtered through 0.22-μm syringe filters (MSI, USA). IAA quantification was carried out by high-performance liquid chromatography (HPLC) using a Shimadzu LC-20AD Prominence system equipped with a UV-VIS detector (SPD-20AV; Shimadzu Corp., Japan). Separation was achieved on a Kinetex C18 reverse-phase column (2.6 µm, 100 Å, 150 × 4.6 mm) at room temperature, using a 20 µl injection loop. The mobile phase consisted of 0.5% acetic acid and methanol (6:4, v/v), with a constant flow rate of 0.6 ml·min⁻¹. Detection was performed at 280 nm, and quantification was done by external calibration using analytical standards.

### In vivo evaluation of P. extremaustralis plant interactions

#### Arabidopsis thaliana assays

*A. thaliana* Col0 (The Arabidopsis Genome Initiative, 2000) seeds were surface sterilized using chlorine gas. The seeds were placed on square plates containing half-strength Murashige & Skoog medium (½ MS) (Murashige and Skoog 1962) and incubated in darkness at 4°C for at least two days to synchronize germination (vernalization). Germination was induced by exposing the seeds to constant light in a vertical position. After 36 h of light exposure, an aliquot of 1.25 µl of an OD^600nm^ =0.05 *P. extremaustralis’* 14-3b suspension, previously grown under aerobic or microaerobic conditions, washed and resuspended in ½ MS medium, was inoculated onto the radicle of each seedling. Controls were performed using ½ MS medium without bacteria. Seedling growth, root length, and root hair length were monitored for 7 days and measured with the ImageJ program.

#### Colonization of A. thaliana

We performed two different colonization assays using mCherry tagged *P. extremaustralis* 14-3b grown either under aerobic or microaerobic conditions. During short term experiments, 5 days old *Arabidopsis* seedlings germinated on ½ MS plates were incubated in a 35 mm Petri dish containing ½ MS and an adjusted bacterial culture (0.1 OD^600nm^).

After 2 h, seedlings were mounted for direct visualization on a confocal microscope. Long term colonization experiments were done as described in section 2.3.1 using GFP tagged strains. Roots and/or cotyledons were observed focusing on different tissue depths for the detection of bacterial cells inside and surrounding plant tissues. In all experiments, the visualization was performed using a confocal microscope (Zeiss LSM900). For short term experiments, mCherry bacteria were visualized with excitation at 560 nm and emission from 560 to 700 nm. For long term invasion assays, we stained plant cell plasma membranes with 1 µM of FM 4-64 dye and excited both FM and GFP at 488 nm. Detection ranged from 480-560 nm for GFP and 560-700 nm for FM4-64.

#### In vivo bacterial AIA production in A. thaliana

DII-VENUS *A. thaliana* seeds (Brunoud et al., 2012), which enable dynamic visualization of auxin distribution through fluorescence in response to auxin signaling, were inoculated as previously described with *P. extremaustralis* 14-3b grown in LB medium under aerobic or microaerobic conditions (DO^600nm^=0.05). Fluorescence levels were quantified at the root tip 96 h after inoculation using a confocal microscope (Zeiss). Root tip stacks were taken with the following parameters: excitation 488 nm, emission 490-620 nm. The resulting pictures were processed with Fiji (Schindelin et al., 2012).

#### Wheat plant assays

Wheat seeds from the commercial cultivars Baguette 601 (NIDERA) and Fuste (DONMARIO) were used. Wheat plants were also grown for 30 days in individual 250 cm^3^ pots containing tyndallized soil as a substrate in a culture chamber with controlled photoperiod (16/8 h light/dark) and temperature (25 ± 5°C). They were inoculated with a bacterial suspension at 10^8^ CFU/g of soil. Controls without bacterial inoculum were performed. Each treatment had 18 independent replications.

#### Bioinformatic and statistical analysis

Bacterial genome was analyzed using programs available online: BLAST (http://blast.ncbi.nlm.nih.gov/Blast.cgi), ClustalW (http://www.ebi.ac.uk/Tools/clustalw/), and other tools included in the RAST server (http://rast.nmpdr.org/) and *Pseudomonas* genome database (http://pseudomonas.com/). Genome sequence accession number is AHIP00000000.1 for *P. extremaustralis* (Tribelli et al., 2012). The significance of each assay was evaluated using the corresponding test using GraphPad Prism software.

## Results

### P. extremaustralis solubilizes phosphorus and produces IAA in vitro and in vivo

In most *Pseudomonas* species, inorganic phosphorus solubilization has been attributed to the secretion of organic acids, with gluconic acid being one of the most important in this process (Iller et al., 2010). Recently, Mayer et al. (2025) performed a genome comparative analysis of genes related to phosphorus solubilization in different *P. extremaustralis* strains.

*P. extremaustralis* 14-3b genome presents *gcd* encoding PQQ-dependent glucose dehydrogenase, converting glucose into gluconic acid, but lacks *gad* coding for gluconate dehydrogenase (that transforms gluconic acid into 2-keto-gluconic acid) present in other *Pseudomonas* species (Miller et al., 2010) We first tested, inorganic phosphorus mobilization in a plate assay using tricalcium phosphate, aluminum phosphate, ferric phosphate, and rock phosphate. Among these, tricalcium phosphate was the only inorganic P source solubilized. No differences were detected using inocula from aerobic or microaerobic cultures (Fig. 1A). We therefore analyzed the organic acid production related to P solubilization in NBRIP-glucose medium by HPLC under aerobic conditions produced by 10^6^ CFU/ml. As expected by the genomic analysis, gluconic acid was detected in *P. extremaustralis,* evidencing the functionality of the gene encoding PQQ-dependent glucose dehydrogenase in this species, as well as pyruvate, with culture media pH value around 3.5-4 after 72 h of incubation (Fig. S1).

**Figure 1.**
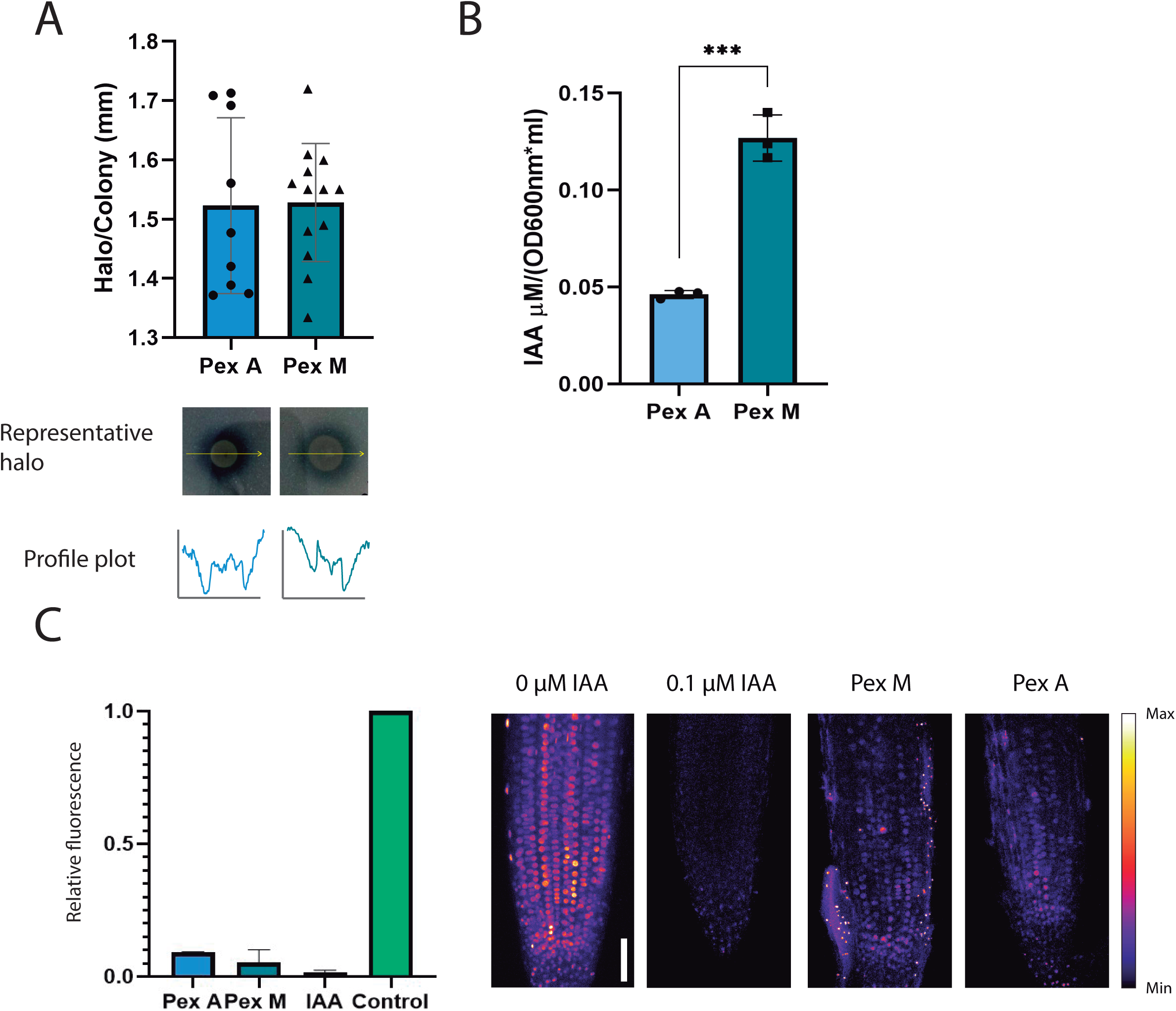
Effect of aeration conditions on production of IAA and phosphorus solubilization in *P. extremaustralis.* A. Calcium phosphate solubilization, representative images of solubilization in NBRIP plates supplemented with tricalcium phosphate and quantification of solubilization halos and their respective intensity plots along the yellow lines. Solubilization is observed as a clear halo surrounding the inoculum. B.Quantification of indole acetic acid production by HPLC under different aeration conditions. Cultures were performed in LB supplemented with tryptophan. Error bars represent the standard deviation of the mean. Asterisks indicate significant differences (P<0.05) using an Unpaired-t test. C: *thaliana* IAA reporter line. Fluorescence of the root of DII-Venus line exposed to 0.1 μM IAA for 30 min or grown in the presence of aerobic (A) or microaerobic (M) cultures of *P. extremaustralis* for 4 days. Scale bar 50 µM. The color scale indicates the range of maximum and minimum signal. Fluorescence is quantified relative to control without IAA addition or inoculum.

Another important metabolite produced by *P. extremaustralis is* the indolic acetic acid (IAA). Regarding IAA, *P. extremaustralis*’ genome possesses the indole-3-acetamide (IAM) pathway, which involves genes encoding the enzyme tryptophan monooxygenase (TMO), which converts L-tryptophan into indole-3-acetamide (IAM), and the indole-3-acetamide hydrolase that produces indol-3-acetic acid (IAA) from IAM. IAA production in tryptophan-supplemented cultures was significantly enhanced under microaerobic conditions in *P. extremaustralis,* showing an average concentration of 0.1269 µM/OD_600_ and 0.04616 µM/OD_600_ under microaerobic and aerobic conditions, respectively (Fig. 1B).

To evaluate in vivo indole-3-acetic acid (IAA) production, we inoculated *Arabidopsis thaliana* seedlings expressing the DII-Venus auxin reporter (Brunoud et al., 2012) with *P. extremaustralis* previously cultured under either aerobic or microaerobic conditions. The DII-VENUS sensor is a genetically encoded tool designed to monitor auxin distribution in vivo, in which the fluorescent protein VENUS fused to domain II of Aux/IAA repressors targets the fusion protein for degradation in response to auxin via the TIR1/AFB receptor pathway. As a result, VENUS fluorescence levels are inversely correlated with local auxin concentrations. Confocal imaging revealed a significant reduction in DII-Venus fluorescence in root cells of inoculated plants, regardless of the bacterial oxygen regime. This decrease in signal reflects elevated IAA levels, as the DII-Venus reporter is degraded in the presence of auxin. In contrast, non-inoculated controls exhibited the strongest fluorescence, consistent with low plant endogenous IAA levels in the observed cells (Fig. 1C).

### P. extremaustralis interacts with Arabidopsis thaliana roots

First, we analyzed the initial binding of a *P. extremaustralis* strain expressing a mCherry protein towards *A. thaliana’s* root. After 2 h, we observed the bacterial cells concentrated along the main root and the root hairs (Fig. 2A). This behavior was observed for both aerobic and microaerobic cultures of *P. extremaustralis*.

**Figure 2.**
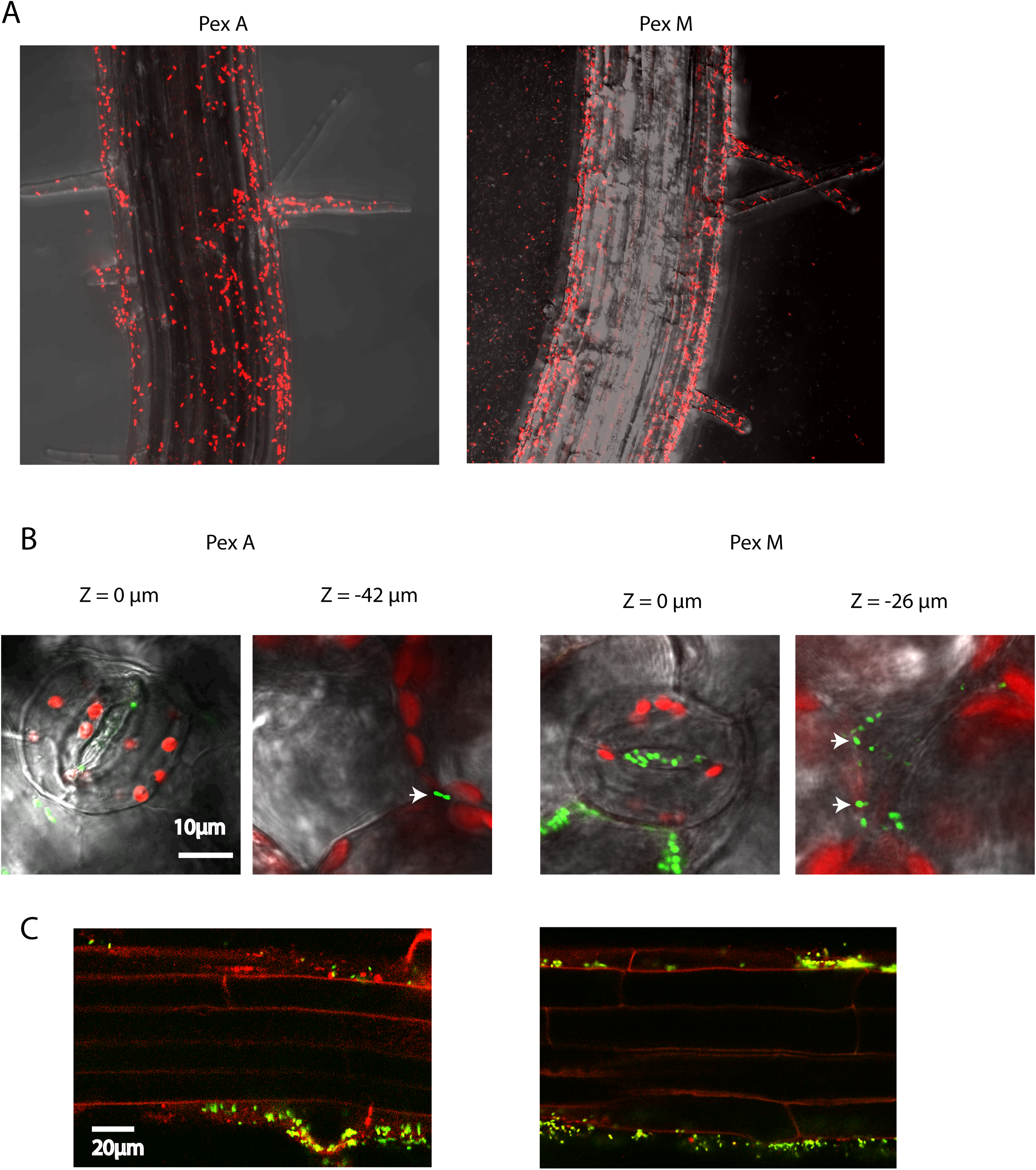
Bacterial initial binding and colonization in *A. thaliana.* A. Confocal images of *P. extremaustralis* expressing mCherry after 2 h of incubation. B-C. Confocal images of plants grown for 4 days after inoculation. B. Images of two different planes of the abaxial leaf side taken at the epidermis surface and in the inner part of the leaf (z values relative to the surface are indicated). Pictures were taken for samples inoculated with aerobic or microaerobic cultures of *P. extremaustralis*. Chlorophyll emission reveals photosynthetic cells inside the leaf. C. Roots from the same samples stained with FM4-64 for plasmatic membrane visualization. Bacteria accumulated close to root hairs. Scale bars: 10 and 20 μm.

Afterwards, we analyzed the *P. extremaustralis* colonization at longer times by using a GFP-expressing strain. After 4 days of inoculation, when looking at the abaxial side of the cotyledons, we observed bacterial cells inside the aerial chamber -in some cases deeper than 40 µm below the stomata surface-, indicating the colonization of the internal part of the leaves (Fig. 2B). Additionally, *P. extremaustraliśs* cells formed a thick biofilm around all root tissues and despite the large number of bacterial cells outside, we didn’t observe any bacteria inside the root tissues in any of the analyzed plants (Fig. 2C). Again, both growth conditions showed the same response regarding the colonization of leaves and roots.

After observing that both aerobic and microaerobically grown bacteria readily colonize *Arabidopsis* plants, we followed their effect on the plant root parameters. The inoculation of *P. extremaustralis* into *A. thaliana* seedlings provoked changes in the root architecture (Fig. 3A). The length of the main root decreased significantly, regardless of the aeration of the cultures from which the inoculum came (Fig. 3B). Lateral roots density (expressed as the number of lateral roots normalized by the length of the main root) was higher in the inoculated plants compared to the control without inoculation but no differences were observed between aeration conditions of the bacterial culture (Fig. 3C). On the other hand, an increase in the length of root hairs was observed in seedlings inoculated with *P. extremaustralis,* compared to the control without inoculum (Fig. 3D). In addition, this parameter also showed significant differences dependent on the aeration of the bacterial culture, with seedlings inoculated with aerobic cultures presenting a longer root hair length (Fig. 3D).

**Figure 3.**
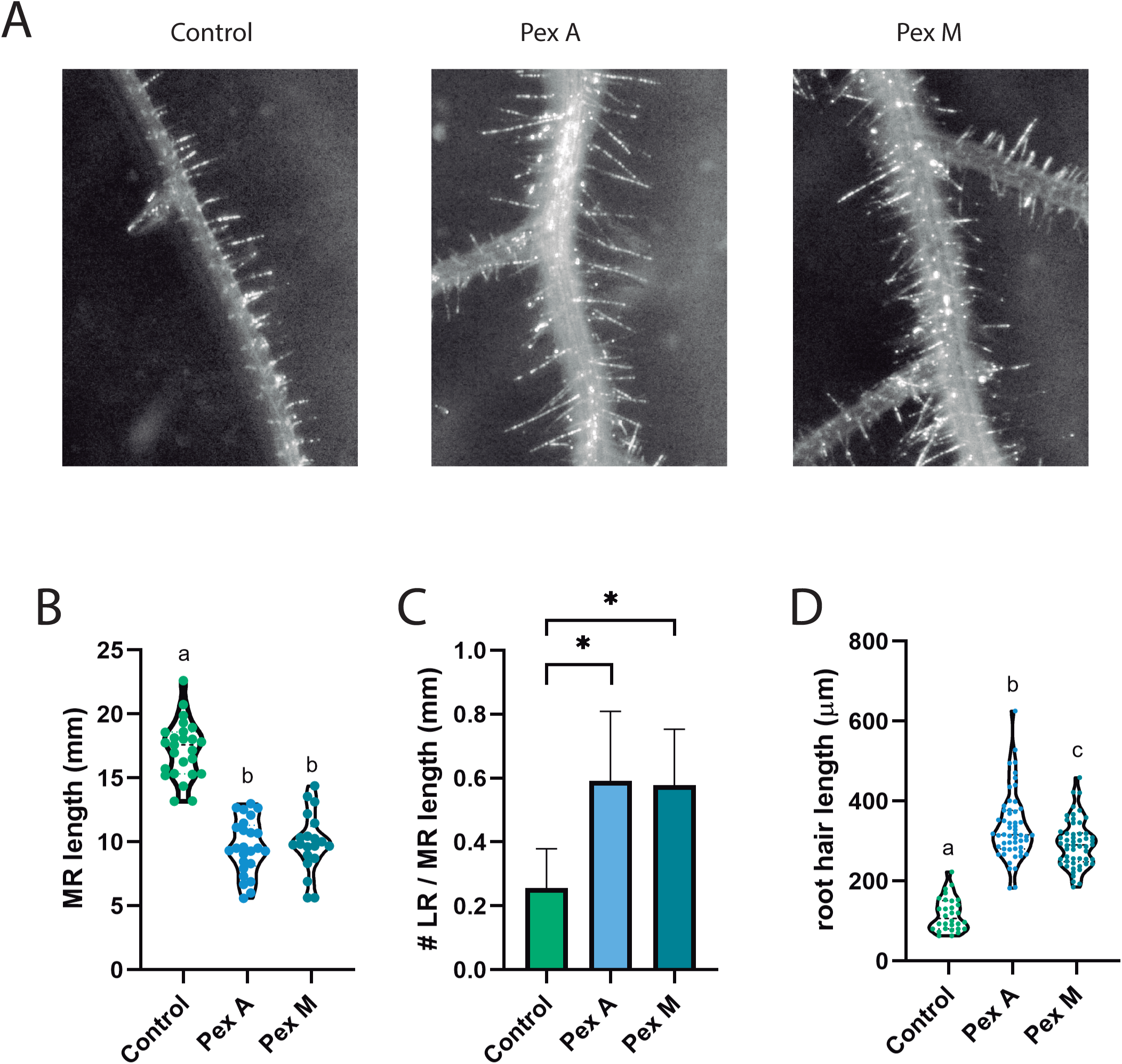
Effect of *P. extremaustralis* grown under different oxygen levels on *A. thaliana* root architecture. A. Representative images of roots. B. Length of main root. C. Root ramification index (LR/MR) D. Comparison of root hair length. Controls without bacteria are also shown. Violin plot displaying data distribution with individual values. Asterisks indicate significant differences (P<0.05) using a One-way Anova with Tukey comparisons.

#### P. extremaustralis inoculation increases steam weight and leaves P content in wheat

As we observed that *P. extremaustralis* can interact satisfactorily with the model plant *A. thaliana*, we aimed to analyze the effect of inoculating a commercial crop, as wheat, with aerobic cultures of this bacterium. Initially, we performed chemotaxis assays using concentrated wheat root exudates. The results obtained showed that *P. extremaustralis* developed a strong response to wheat exudates after 15 min of incubation, visualized as a white ring surrounding the center of the plate when root exudates were added (Fig. S2).

To evaluate the effect of *P. extremaustralis* on wheat plant nutrition and health, a trial was conducted in which wheat plants were grown in pots containing tyndallized soil containing 1.53 g/kg total N, 14.90 g/kg total C, 13 mg/kg of assimilable phosphorus content, and pH 5.1. *P. extremaustralis* cultured aerobically was inoculated at 10^8^ CFU/g of soil, and plants were grown for 30 days. After this period, the stem dry weight was significantly higher in inoculated plants compared with the control (Unpaired student t test; P=0.0107), while the root dry weight was similar between inoculated and non-inoculated plants (Fig. 4A and B). No differences were observed in total chlorophyll content in leaves, while total phosphorus content in this tissue was higher in inoculated plants (Unpaired student t-test; P=0.0242), showing the impact of *P. extremaustralis* interaction (Fig. 4C and D). These trials showed a beneficial bacterial effect in the aboveground parts of the plants, reflected in higher P content in leaves and higher stem development.

**Figure 4.**
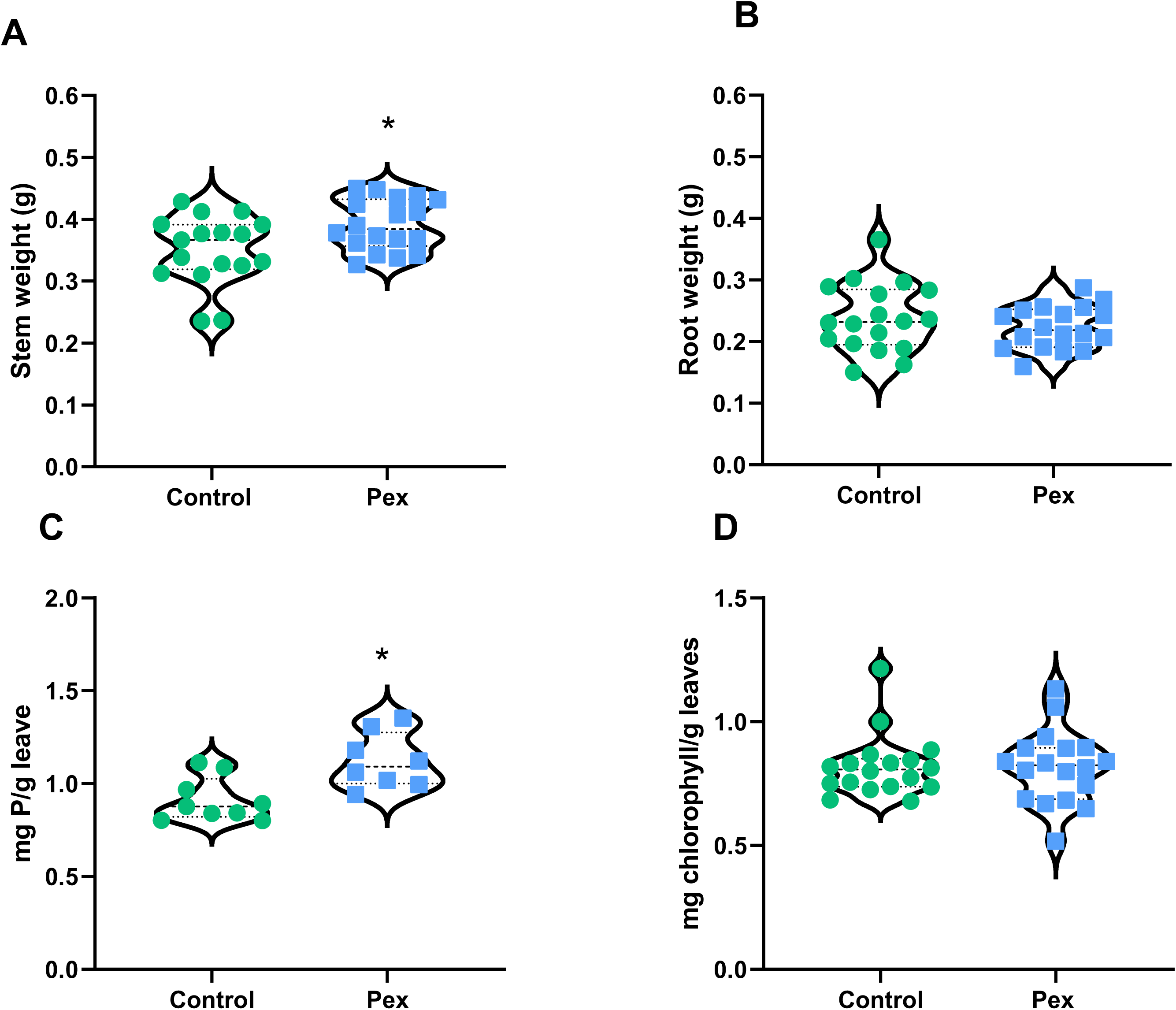
Growth parameters of 30-day-old wheat plants. Plants were grown in 250 cm^3^ pots with tyndallized soil, inoculated with *P. extremaustralis* and uninoculated. A. Stem dry weight (N=18); B. Root dry weight (N=18); C.: Total phosphorus in leaves (N=9); D. Chlorophyll content in leaves (N=18). Violin plot displaying data distribution with individual values. Asterisks indicate significant differences using an unpaired t-test (P<0.05).

## Discussion

The intensification of agriculture has led to widespread misuse of soils and excessive dependence on synthetic fertilizers, contributing to nutrient depletion and long-term soil degradation. This unsustainable model undermines both crop productivity and environmental integrity. Microbial bioinoculants, more particularly the plant growth promoter bacteria (PGPB), represent a viable strategy to enhance nutrient cycling and soil health while minimizing chemical input. Their application is increasingly recognized as a key component of sustainable agricultural systems.

The PGPR characteristics of *P. extremaustralis* were studied using genomic and traditional microbiology tools in vitro and also in vivo by analyzing the interaction with plants. This species shows phylogenetic closeness to *Pseudomonas* species related to plant growth promotion, can tolerate different types of stress, and produces polyhydroxyalkanoates; in addition, knowledge of its genome showed several interesting traits (Raiger Iustman et al., 2015).

To verify the *P. extremaustralis’s* interaction with plants, we carried out different tests using both the plant model *A. thaliana* and a commercial crop, wheat. Chemotaxis is a relevant characteristic in the plant-bacteria relationship involving different regulator pathways and cellular structures like flagella. In this work, *P. extremaustralis* was attracted immediately by root exudates of wheat plants in in vitro experiments and was attracted to *A. thaliana* roots, as was shown by confocal microscopy. This process is recognized as the first step in the plant-bacteria interaction (Knights et al., 2016). Feng et al. (2022) have reviewed the different chemoeffectors present in the root exudates, being aminoacids and organic acids the most relevant, but the composition is also affected by stressful conditions (Chai and Scachtman, 2022; Wang et al., 2024). *P. fluorescens* Pf0-1 senses a broad range of amino and organic acids through specific chemoreceptors: CtaB detects most amino acids, including glutamine and glutamic acid; CtaC responds to a subset including arginine, cystine, and methionine; and McpT mediates chemotaxis toward L-malate and succinate, and other aminoacids and organic compounds may be sensed via yet unidentified mechanisms (Oku et al., 2012). *P. extremaustralis* has the complete chemotactic systems that encode the Che2 and Che3 proteins, which are absent in other PGPB bacteria like *P. protegens* Pf-5. The differences in these systems could be related to the different behavior towards root exudates. Another important feature for the initial plant-bacteria interaction is flagellar motility (Capdevila et al., 2004; de Weert et al. 2002). In most of the *Pseudomonas*, the genes for the synthesis of polar flagella are distributed in three groups (Redondo-Nieto et al. 2013). *P. extremaustralis* also has another cluster that has 40 open reading frames that encode a complete flagellar system, only described in *P. fluorescens* F113, which is characterized by its high competence in the rhizosphere (Barahona et al., 2016).

Positive chemotaxis towards the plant root is the first step in the colonization process, but to achieve a successful colonization, the development of stable interactions is essential, which can be observed as the development of bacterial colonies on the root surface or the establishment in the periplasmic and intercellular spaces, among others (Knights et al., 2016). The distinct zonation of plant roots into the root tip, elongation zone, and maturation zone (Verbon and Liberman 2016) is fundamental to plant-bacteria interactions and notably influences bacterial colonization. Unge and Jansson (2001) reported the colonization of wheat roots by *P. fluorescens* SBW25 at 6 days after inoculation. Moreover, *P. fluorescens* F113 inoculated in seeds was detected in different tissues such as roots and leaves, though colony counting, but no specific localization within the leaves was described (Lally et al., 2017). In this work, we showed that in *A. thaliana,* bacterial cells were present in cotyledons and root surfaces. Bacteria were also present at the aerial chamber below leaf stomata but not inside the roots, although they were surrounding the rotor and lateral hairs, providing new experimental data of the substomatal cavity localization of a bacterium with PGPB characteristics like *P. extremaustralis*.

It is well known that rizobacteria with PGP characteristics act through direct and indirect mechanisms, including phosphorus (P) solubilization, indole acetic acid biosynthesis (IAA), siderophores and volatile compounds production. Soil phosphorus (P) deficiency is a major worldwide constraint to crop yield, and due to phosphorus sorption of most soils, less than 20 % of applied fertilizer P may be up taken by plants during the growing season (Shen et al., 2018). Due to its low mobility in soil, phosphate reaches the root surface primarily by diffusion, although its limited diffusion coefficient can be enhanced by increasing its concentration in the soil solution (Lambers et al. 2008). Dense root branching reduces the diffusion distance and improves phosphate interception at the root surface (Lambers et al. 2006; Postma et al. 2014), and the use of free-living bacteria capable of solubilizing P increases its potential absorption by the crop. The presence of genes related to phosphorus solubilization was reported in different strains of *P. extremaustralis* by Mayer et al. (2025). In this work, we showed the inorganic phosphorus solubilization of cultures grown previously under different oxygen conditions and determined the production of gluconic acid, confirming experimentally the *P. extremaustralis* capability of phosphorus solubilization. In pots, wheat showed an increment in P in leaves, showing that *P. extremaustralis* not only possessed an in-vitro P solubilization activity but also produced a difference in the nutritional condition of wheat. Interestingly, the role of phosphorus content in tolerance to abiotic stresses such as heat, drought, salinity, waterlogging, elevated CO₂, and heavy metals was revised by Lamberts et al., and it was even reported that the P content can alter root characteristics and stomata features (Lamberts et al., 2022).

As we described above, the root is the main organ in contact with bacteria from the rhizosphere, and the crosstalk between the bacteria and the plant involves not only the root exudates but also the production and secretion of bioactive compounds by the bacteria (Massalha et al., 2017). We demonstrated the *in vitro* IAA production of *P. extremaustralis* cultures under both aerobic and microaerobic being higher under microaerobic conditions. Moreover, our experiments demonstrated that *A. thaliana i*s sensing the IAA produced by *P. extremaustralis*, showing differences between non-inoculated plants and those inoculated, although no differences were observed between the oxygen conditions in which the bacterium was grown. The root architecture was strongly modified when *A. thaliana* was inoculated with *P. extremaustralis,* showing a shorter principal root but a significant increment in the root hair length. Environmental stresses like nutrient and water scarcity severely impact plant productivity, a challenge set to worsen with climate change. Consequently, root hairs are a critical target for developing stress-resilient crops, as they enhance a plant’s ability to acquire soil resources due to the increment of root surface area and the root-soil contact (White & Kirkegaard, 2010; Zhang et al., 2018; Marin et al., 2021; Duddek et al., 2022). This expanded reach, coupled with their ability to access finer soil pores, notably improves nutrient and water uptake, particularly for immobile nutrients like phosphorus as we described above (Keyes et al., 2013).

As mentioned, *P. extremaustralis* 14-3b was isolated from a water sample from a temporary pond in Antarctica. Considering the site of isolation, Punta Cierva on the Danco coast, an area of scientific interest on the west of the Antarctic Peninsula (Ayub et al., 2004), the presence of PGP characteristics in its genome seems strange. However, this environment has some peculiarities, such as a benign microclimate, considering the latitude, compared to other regions located further north in this continent, an average monthly temperature that ranges between 1.8-2.2° C, the good develop of vegetation, with cover of mosses, grasses, and lichens during summer, when the thaw occurs, and the presence of many bird species (Quintana et al. 2000, Quintana 2001). The geomorphological, climatic, and biological characteristics provide many habitats, suggesting that *P. extremaustralis* could also be in contact with vegetation in that environment. In addition, subsequently, different strains of *P. extremaustralis* were isolated from different environments, including soils. Recently, Mayer et al. (2025) analyzed the presence of genetic determinants related to phosphate solubilization in all the available genomes of *P. extremaustralis* strains from different environments, demonstrating that this capacity is widespread in this species. Moreover, their results also showed the *in vitro* ability to solubilize inorganic P of two *P. extremaustralis* strains isolated from pisciculture sludge residues, indicating the role in phosphorus-cycling and highlighting the potential of this species as a facilitator of plant-available phosphorus, which could be interesting for the development of sustainable agriculture.

In summary, in this study, we provide comprehensive evidence that *P. extremaustralis* possesses multiple traits associated with plant growth-promoting bacteria, including chemotactic attraction to root exudates, colonization of aerial and root tissues, phosphorus solubilization, and IAA production. Notably, we employed an in vivo biosensor-based approach to confirm that the IAA synthesized by *P. extremaustralis* is biologically active and perceived by *A. thaliana*, leading to marked changes in root architecture. These findings reinforce the potential of *P. extremaustralis* as a functional bioinoculant.

## Supplementary Figures

**Supplementary Figure 1.** Organic acid production. Time course of gluconic acid and pyruvic acid production under aerobic conditions in NBRIP broth supplemented with tricalcium phosphate quantified by HPLC. Values represent media and SD of three independent experiments.

**Supplementary Figure 2.** Chemotactic response of *P. extremaustralis* towards wheat root exudates. A. Homogeneous suspension of bacteria (DO600nm=0.8) in chemotaxis buffer. B. 10 µl of concentrated exudates were added in the center of the plate.

## Notes

### Competing Interest Statement

The authors have declared no competing interest.

### Funding

- University of Buenos Aires: 20020220400198BA

