## Supplementary figures and images for "Functional auxin signaling and phosphorus acquisition triggered in planta by the extremophile bacterium *Pseudomonas extremaustralis*"

### Supplementary Figures

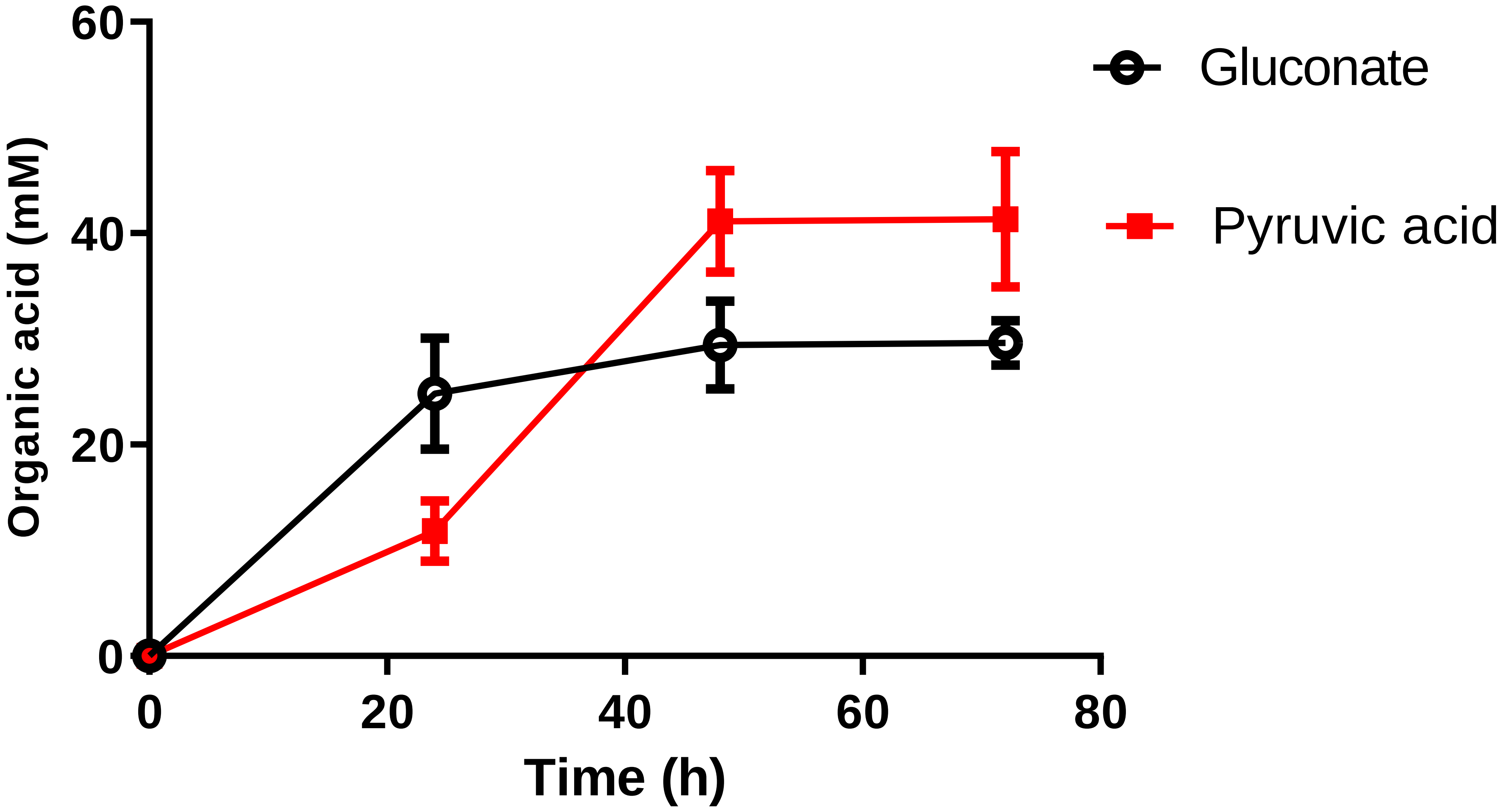

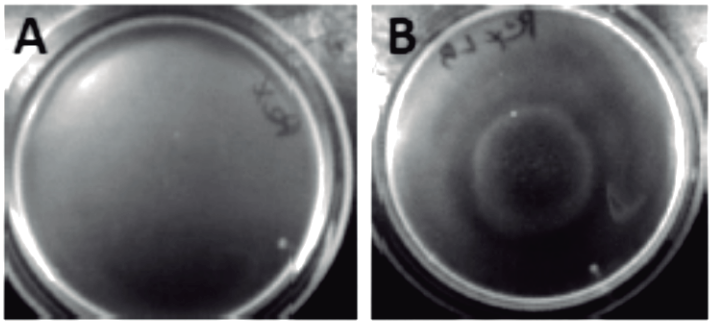


Pex suspension

Pex suspension
+
Root exudates
